# Euo is Essential for Transcriptional Priming of *Chlamydia trachomatis* Elementary Bodies to Facilitate Secondary Infection

**DOI:** 10.64898/2026.08.22.746413

**Authors:** Cody Appa, Nicole A. Grieshaber, Colleen C. Monahan, Connor D. Blum, Anders Omsland, Scott S. Grieshaber

**Affiliations:** Department of Biological Sciences, University of Idaho, Moscow, ID 83844; Paul G. Allen School for Global Health, Washington State University, Pullman, WA 99164

**Author notes:** Address correspondence to: Scott S. Grieshaber, Department of Biological Sciences, University of Idaho, Moscow, ID 83844.

## Abstract

The phylum *Chlamydiota* comprises obligate intracellular bacteria characterized by a highly conserved, biphasic developmental cycle. This cycle involves the transition between the infectious, metabolically quiescent elementary body (EB) and the non-infectious, replicative reticulate body (RB). While the morphological transitions of the developmental cycle are well-documented, the regulatory mechanisms governing these phenotypic shifts remain poorly understood. A primary candidate for this regulation is Euo, a conserved, phylum-specific helix- loop-helix transcription factor hypothesized to repress late-cycle genes and prevent premature differentiation.

In this study, we employed CRISPR interference (CRISPRi) to knockdown *euo* expression in *Chlamydia trachomatis* to further elucidate its role in developmental regulation. Unexpectedly, *euo* knockdown did not significantly disrupt the primary developmental cycle; progression through RB replication, the formation of intermediate bodies (IBs), and the kinetics of late-gene expression remained largely comparable to wild-type. However, we observed a significant reduction in the production of infectious progeny.

Detailed analysis revealed that while EBs were still produced and capable of entering host cells after knock down of *euo*, these EBs exhibited dysregulated gene expression during the germination phase of a new infection cycle. Consequently, these bacteria failed to establish a productive secondary infection. These results suggest that rather than acting as a developmental switch for differentiation during the initial infection, Euo is essential for the proper programming of EBs, ensuring transcriptional competence upon re-infection of a host cell.

## Introduction

The phylum *Chlamydiota* consists of obligate intracellular bacterial pathogens/endosymbionts of eukaryotic organisms. These bacteria undergo a developmental cycle producing phenotypically distinct and specialized cell forms responsible for either amplification or dissemination. The *Chlamydiota* contains important pathogens of humans and vertebrate animals. In humans, *Chlamydia. pneumoniae* causes respiratory infections including pneumonia, while *Chlamydia trachomatis* (Ctr) infections cause blinding trachoma, urethritis in men and cervicitis in women. Although the bacteria in this phylum infect eukaryotic cells across the kingdom, the broad strokes of the developmental cycle are highly conserved.

This complex intracellular developmental cycle consists of multiple cell forms; the elementary body (EB), the reticulate body (RB) and the intermediate body (IB) (1, 2), and has mostly been characterized for the human pathogen Ctr. The Ctr EB initiates infection of host cells via a type III secretion system (T3SS) and pre-formed effectors (3–5) that promote pathogen phagocytosis and entry into the targeted cell. After entry the developmental cycle follows a complex cell phenotype progression. The endocytosed EB initiates immediate early gene expression (0-4 hpi) that controls the exit of the chlamydial inclusion from the endocytic pathway and initiates dynein dependent trafficking to the host cell microtubule organizing center (MTOC) (6, 7), ultimately interacting intimately with the Golgi and ER (8, 9). After trafficking the EB completes germination and becomes replication competent at ∼10 hpi (Ctr serovar L2) (10). The RBs undergo several rounds of replication before generating IBs, which subsequently mature into infectious EBs over an 8-to-10-hour window (11, 12). The mature RBs continue to produce IBs, acting as a mother cell population (11). The factors that regulate the developmental cycle are currently poorly understood.

The chlamydial genome encodes a number of DNA binding proteins that are conserved across the phylum but are not found in other bacterial lineages (13). One such protein is the Early Upstream Open Reading Frame (Euo) protein. Euo is a helix-loop-helix containing transcription factor and has been proposed to act as a regulator of cell form differentiation (14–17). Euo is expressed in the RB very early upon infection and has been shown to bind to promoter regions of several late genes, potentially repressing IB and EB gene expression (14, 15, 17). Ectopic expression of Euo results in reversible arrest of the development cycle (16, 18). The arrested chlamydial cells are trapped phenotypically at an early IB stage which is dependent on continued expression of Euo (16). The developmental cycle resumes with wild-type kinetics when the inducer is removed and ectopic expression ceases (16).

In this study, we used CRISPRi to knockdown *euo* expression. Knockdown of *euo* significantly reduced the production of infectious progeny. Surprisingly, this knockdown had little effect on the primary developmental cycle as measured by progression through RB replication, IB production and EB gene expression kinetics and morphological imaging studies. However, upon secondary infection, the EBs entered host cells with apparent dysregulated gene expression during germination, and ultimately failed to produce a productive infection.

## Results

### Knockdown of *euo* leads to a decrease in the production of infectious progeny

We previously demonstrated that overexpression of Euo inhibited division and maturation in the early IB without impacting the RB or the committed step to IB production (16). To further understand the impact of Euo on chlamydial development, we utilized a CRISPR-dCas12 gene knockdown system (CRISPRi) with a guide RNA targeting the *euo* gene to perform targeted knockdown under the control of an inducible promoter. We generated two *euo* knockdown strains, L2-BsciDng-dCas12-*euo*g and L2-PsciDng-dCas12-*euo*g (Fig. S1). The dual fluorescent reporter cassettes of each strain expressed mNeonGreen driven by the RB-specific *incD* promoter (Dng), and either Scarlet-I driven by the EB-specific *hctB* promoter (Bsci) or Scarlet-I driven by the IB specific promoter porB (Psci) (18).

To verify that *euo* mRNA expression was knocked down, monolayers of Cos-7 cells were infected with L2-BsciDng-dCas12-*euo*g strain at an MOI of ∼0.3, and induced for dCas12 expression. At 16 hpi, *euo* transcripts levels were evaluated by fluorescent in situ hybridization (FISH) and confocal microscopy (Fig. 1A). Induced and uninduced samples displayed morphologically similar inclusions containing *incD* promoter–positive RBs. However, *euo* transcripts were readily detectable only in the uninduced samples, indicating efficient CRISPRi-mediated knockdown upon induction (Fig. 1A). To assess the functional impact on development, we quantified progeny EBs by harvesting infectious particles from induced and uninduced cultures and measured inclusion-forming units (IFU) in a reinfection assay. *euo* knockdown resulted in a marked reduction in IFU compared with uninduced controls (Fig. 1B).

**Figure. 1:**
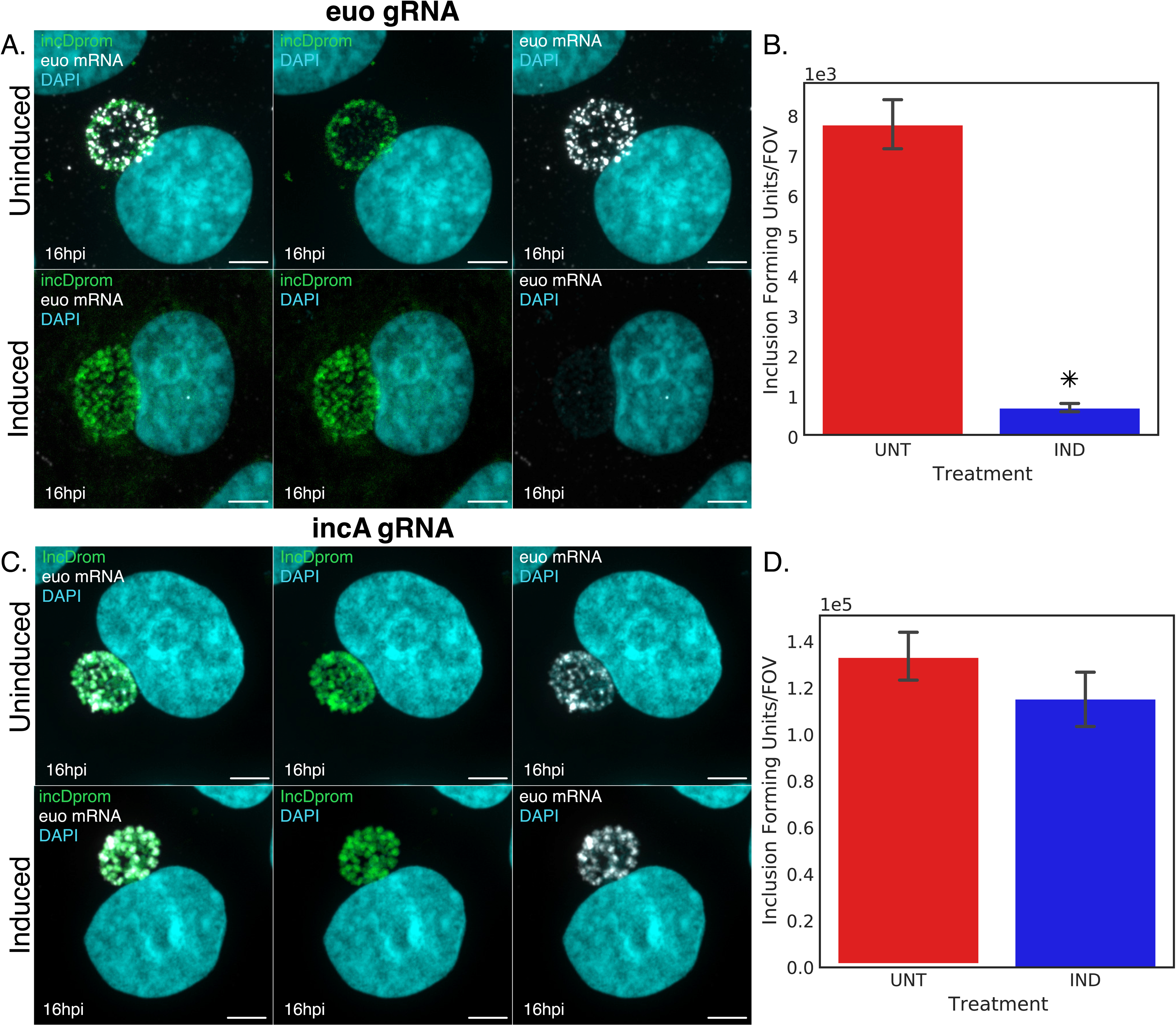
Knockdown of *euo* expression. A) Representative confocal micrographs of Cos-7 monolayers infected with L2-BsciDng-dCas12-*euo*g. Cultures were either left uninduced or induced with 3 ng/mL anhydrotetracycline (aTc) at 0 hpi. The RB-associated *incD* promoter activity is shown in green, and *euo* mRNA transcripts, detected by FISH, are shown in white. DNA was counterstained with DAPI (blue). Samples were fixed at 16 hpi; MOI ≈ 0.3. B) Progeny infectivity assay (IFU). Secondary monolayers were infected with purified L2-BsciDng-dCas12-*euo*g EBs harvested from the *euo* knockdown primary infection at 48 hpi. Inocula were normalized to chlamydial genome counts (ddPCR). Inclusion formation was quantified via automated microscopy. C) Confocal micrographs of the *incA* control strain (L2-BsciEng-dCas12-*incA*g) under uninduced or induced conditions. The *euo* promoter signal is shown in green, and *euo* mRNA transcripts are shown in white, confirming that *incA* knockdown does not affect *euo* expression. D) Progeny infectivity assay for L2-BsciEng-dCas12-*incA*g, performed as described in (B). No significant reduction in IFU was observed upon *incA* knockdown. Scale bar = 15 µm. Data represent the mean ± SD of three independent experiments. * = p < 0.01.

We next examined the control strain L2-BsciDng-dCas12-*incA*g, which targets *incA*, a gene that has been shown to be dispensable for normal progression of the *C. trachomatis* developmental cycle (19, 20). Inclusions from induced and uninduced cultures were indistinguishable and both displayed robust *euo* FISH signal (Fig. 1C). Consistent with expectations, no significant difference in EB production was observed between induced and uninduced conditions for the control strain (Fig. 1D).

### Knockdown of *euo* does not alter the developmental kinetics of the primary infection

To determine whether Euo depletion affects the timing or progression of the chlamydial developmental cycle, we monitored reporter expression dynamics for both L2-BsciDng-dCas12- *euo*g and L2-PsciDng-dCas12-*euo*g. Using single-inclusion live-cell imaging, the reporter cassettes allow for quantitative tracking of RB cell number expansion, IB production, and EB differentiation over the course of infection (11, 12).

Host cell monolayers were infected with each knockdown strain and treated with either aTc to induce *euo* knockdown or vehicle control at infection. Infected monolayers were imaged in both green and red channels every 30 minutes for 48 hours, and promoter-specific fluorescence intensities were measured for individual inclusions. Average expression profiles were then plotted to assess developmental progression (Fig. 2A).

**Figure 2:**
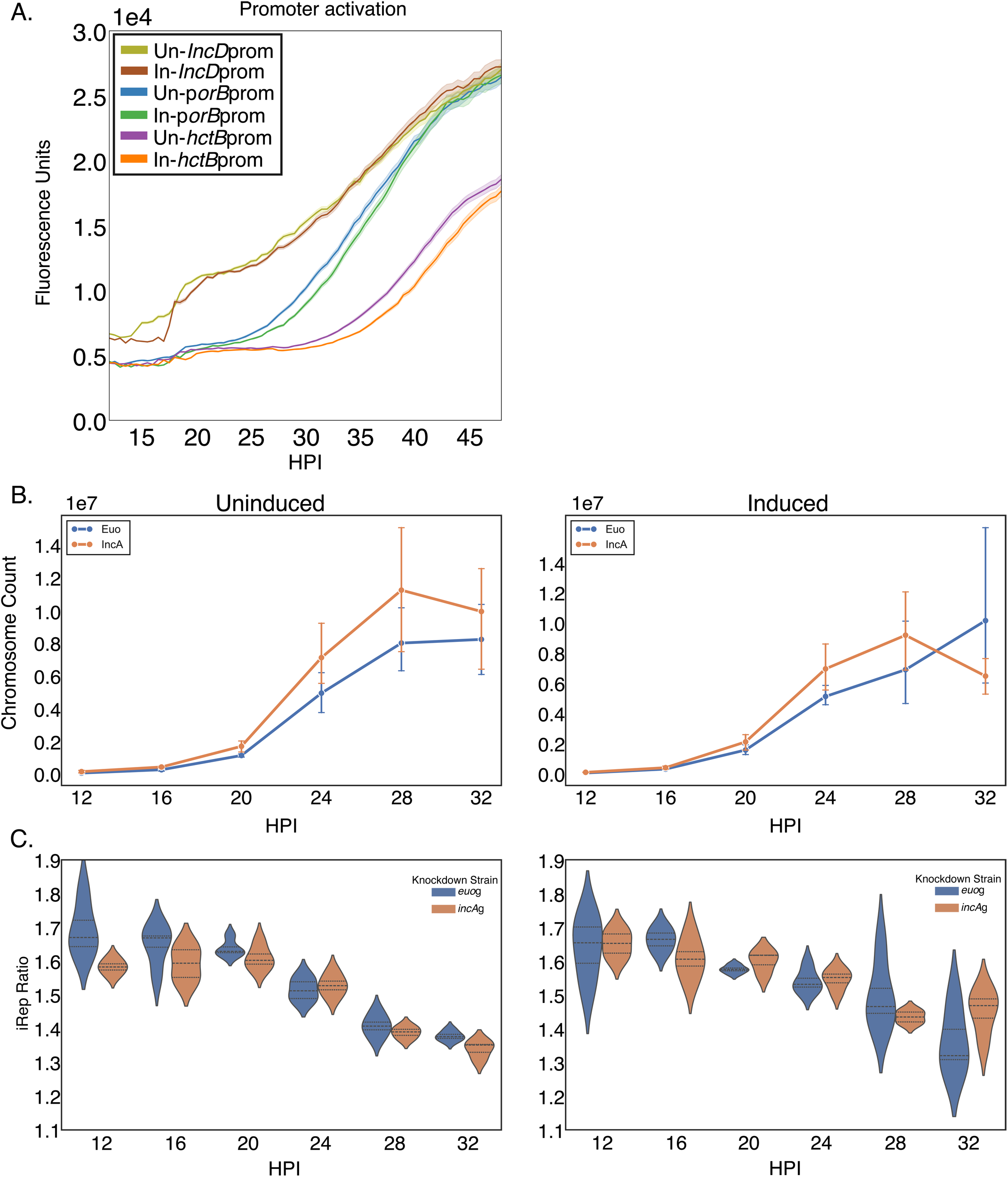
Knockdown of *euo* does not alter primary developmental kinetics or chromosomal replication. A) Temporal expression kinetics via live-cell fluorescence tracking of *Chlamydia* developmental transitions. Monolayers were infected with L2-PsciDng-dCas12-*euo*g under either uninduced or induced (3 ng/mL aTc at 0 hpi) conditions and imaged every 30 minutes for 50 hours. Lines represent the normalized fluorescence intensity of stage-specific reporters: *incD*prom (RB-specific; yellow and red), *porB*prom (IB-specific; blue and green), and *hctB*prom (EB-specific; purple and orange). No significant temporal shift was observed between induced and uninduced samples. B) Growth curves showing accumulation of chlamydial chromosomes over time in uninduced or induced cultures of L2-PsciDng-dCas12-*euo*g and the control strain L2-PsciEng-dCas12*-incA*g, quantified by ddPCR. C) Analysis of replication dynamics by measuring the Index of Replication (iRep) values calculated from ddPCR data for the indicated strains between 12 and 32 hpi. Error bars represent SEM; n>100 droplets per treatment for ddPCR measurements.

Induction of *euo* knockdown had no detectable impact on the overall timing or sequence of developmental transitions. In both induced and uninduced samples, *incD*prom expression was observed first, marking RB-dominant early development, followed by activation of the *porB*prom reporter corresponding to appearance of IBs, and finally *hctB*prom activation consistent with EB maturation (Fig. 2A). These data indicate that *euo* knockdown does not impede the primary developmental kinetics of the infection.

### Euo knockdown does not alter chlamydial genome replication dynamics

We next assessed whether *euo* knockdown affected chromosomal replication during the developmental cycle. Cos-7 monolayers were infected with L2-BsciDng-dCas12-*euo*g in the presence or absence of aTc. The L2-BsciDng-dCas12-*incA*g strain served as a gene-specific knockdown control. Consistent with our live-cell imaging results, *euo* knockdown did not affect the overall accumulation of chlamydial chromosomes. Infections exhibited indistinguishable increases in genome copy number over time regardless of *euo* expression, a pattern also observed for the *incA* control strain (Fig. 2B).

We previously demonstrated that the index of replication (iRep) is a sensitive method for monitoring chlamydial developmental progression, as it captures the ratio of chromosomal coverage at the origin (ori) relative to the terminus (term), thereby reflecting replication initiation frequency (15, 20). During early infection, when RBs predominate and are actively replicating, the iRep is typically ∼1.5–2. As EBs accumulate later in development and cease DNA replication, the population-level iRep declines toward 1 (16, 19).

To determine whether *euo* knockdown impacted the regulation of chromosomal replication, we used ddPCR to measure iRep across the developmental cycle in induced and uninduced cultures of both the *euo*g and *incA*g strains. In uninduced infections, iRep values at 12 hpi, when the population is dominated by replicating RBs, were approximately 1.7 for both strains (Fig. 2C). Values remained >1.5 until ∼24 hpi, after which iRep declined below 1.4 between 24– 33 hpi, coinciding with the onset of EB accumulation. Induction of *euo* knockdown produced an identical temporal pattern, with no measurable shift in the timing or magnitude of iRep changes, as compared to the uninduced control (Fig. 2C). The only notable difference was increased variability in iRep measurements during the EB-formation phase in the *euo* knockdown strain, although the overall trend remained comparable to controls.

### Knockdown of *euo* altered the spatial organization of the IB cell but did not affect the morphology of individual chlamydial cell forms

Because *euo* knockdown markedly reduced infectious progeny without affecting genome replication or the temporal progression of cell_form–specific gene expression, we next examined its impact on IB and EB phenotypes. To visualize IBs during development, host cells were infected with *C. trachomatis* L2-PsciDng-dCas12-*euo*g, in which the *porB* promoter (Psci) drives scarlet-I expression in IB and EB forms, while the *incD* promoter drives neongreen expression in RBs only. Infected host cell monolayers were either induced or left uninduced for dCas12-mediated knockdown and fixed at 30 hpi. DNA was visualized by DAPI staining (Fig. 3A).

**Figure 3:**
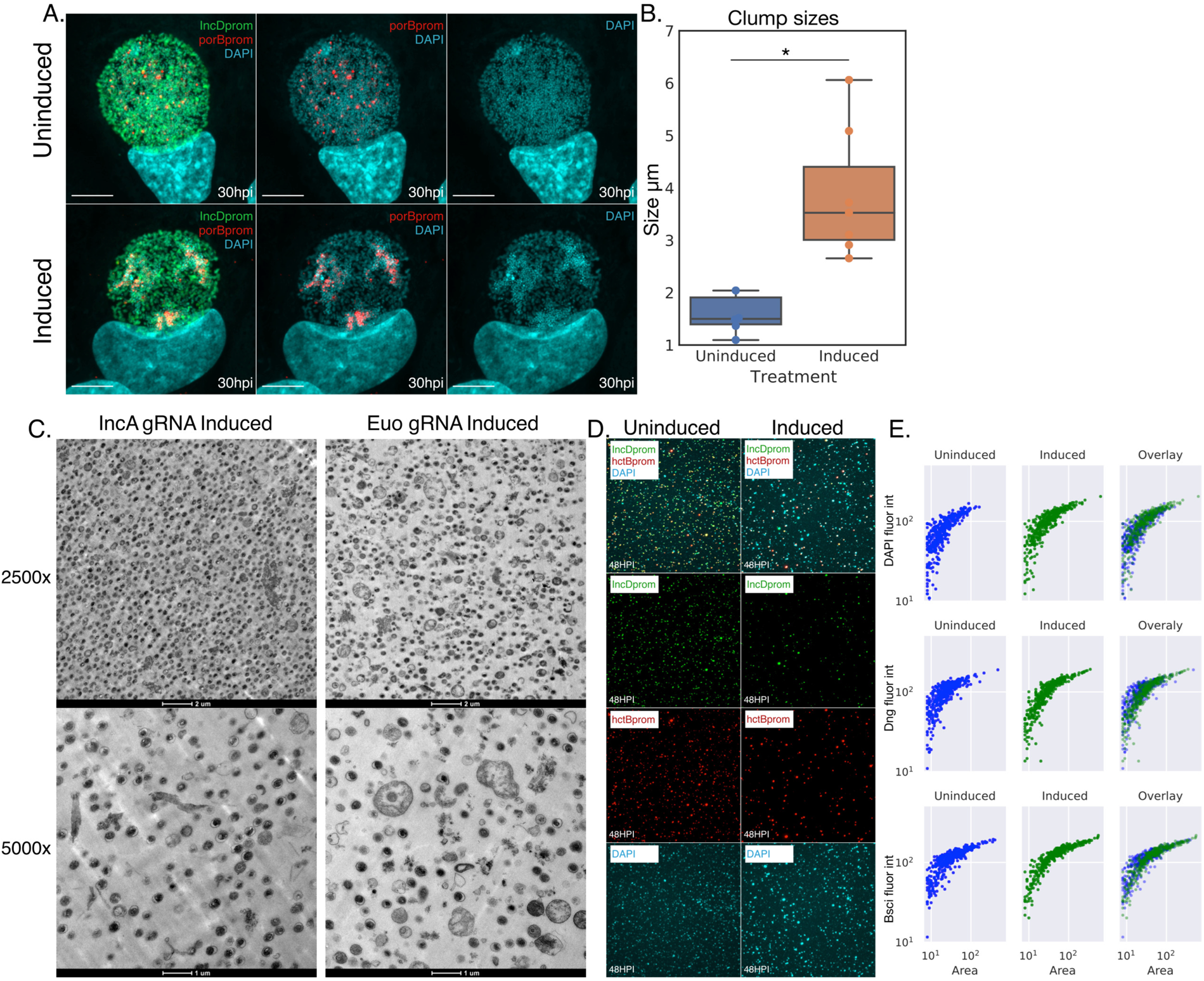
Knockdown of *euo* alters IB spatial organization but does not impact individual cell morphology. A) Spatial organization of developmental forms was assessed by confocal microscopy of monolayers infected with L2-PsciDng-dCas12-*euo*g, either uninduced or induced with 3 ng/mL aTc at 0 hpi. Samples were fixed at 30 hpi. The *incD*prom (RB) signal is shown in green and the *porB*prom (IB/EB) signal in red. B) Quantification of IB aggregation was determined via measurement of the mean diameter (μm) of *porB*prom-positive structures within inclusions. *euo* knockdown resulted in a significant shift from individual IBs to larger “IB clumps” or aggregates. C) Ultrastructural analysis via transmission electron micrographs (TEM) of EBs purified from L2-BsciDng-dCas12-*euo*g infections (uninduced or induced) harvested at 48 hpi. Images were acquired at 2500x and 5000x magnification, showing typical nucleoid condensation and electron-dense morphology in both conditions. D) Morphology of purified progeny. Confocal micrographs of isolated chlamydial forms harvested by sonication at 48 hpi and immobilized on poly-L-lysine-coated slides. RBs (*incD*prom+) are labeled in green and EBs (*hctB*prom+) in red; DNA is counterstained with DAPI (blue). E) Quantification of cell size (area in µm) for total chlamydial forms (DAPI+), RBs (*incD*prom+), and EBs (*hctB*prom+) using ImageJ-based segmentation plotted against fluorescent signal intensity. No significant differences in individual cell size were observed between uninduced and *euo* knockdown conditions.

Consistent with earlier observations, *euo* knockdown did not noticeably affect the distribution or abundance of *incD*prom+ RBs. In contrast, *porB*prom+ bacteria displayed a pronounced alteration in spatial organization, forming discrete clusters within inclusions rather than appearing as individual or dividing cells.

To quantify this phenotype, we measured the average size of resolvable *porB*prom+ IB structures from 10 inclusions per condition using automated thresholding in ImageJ (Fig. 3B). In uninduced samples, most *porB*prom+ structures measured 1.5–2 µm in diameter, consistent with single IBs or dividing cells. In contrast, following *euo* knockdown, *porB*prom+ clusters were substantially larger, with the majority measuring 3–4.5 µm in diameter and some reaching up to 6 µm, indicative of aberrant aggregation of the IB cell form.

To assess whether *euo* knockdown resulted in gross morphological defects in chlamydial developmental forms, we next analyzed purified EBs by TEM and confocal microscopy. For TEM, cells were infected with either L2-BsciDng-dCas12-*euo*g or L2-BsciDng-dCas12-*incA*g as a control and induced for dCas12 expression. EBs were harvested at 48 hpi and fixed. Micrographs revealed a nearly homogeneous population of electron-dense EBs in both the *euo*g and *incA*g samples, with no obvious ultrastructural differences between conditions (Fig. 3C).

For confocal analysis, Cos-7 cells were infected with L2-BsciDng-dCas12-*euo*g under induced or uninduced conditions. At approximately 48 hpi, chlamydial forms were isolated. The chlamydial cells were fixed onto coverslips and imaged by confocal microscopy. Both conditions displayed similar populations, consisting predominantly of small red cells (i.e., EBs) and fewer larger green cells (i.e., RBs) (Fig. 3D).

Using ImageJ-based segmentation, we quantified cell size and fluorescence intensity for *incD*prom, *hctB*prom, and DAPI signals and plotted signal intensity as a function of cell size (Fig. 3E). No discernible differences were observed between induced and uninduced samples for any cell form or fluorescence marker, indicating that *euo* knockdown does not grossly alter individual chlamydial cell morphology or cell_form–specific marker expression.

The observed changes in chlamydial cell organization and distribution within the inclusion was reminiscent of the inclusion phenotypes of Ctr strains with defects in glycogen synthesis (20–23). To investigate whether the morphological alterations correlated with disrupted glycogen storage, inclusions were assessed for polysaccharide accumulation using Periodic Acid-Schiff (PAS) staining (24, 25) at 45 hours post-infection (hpi). Cos-7 cells were infected with Ctr L2 wild-type (wt), the plasmidless strain L2R, and the inducible knockdown strains L2-BsciDng-dCas12-*euo*g or L2-BsciDng-dCas12-*incA*g under both induced and uninduced conditions. As anticipated, L2 wt inclusions exhibited robust, intense PAS staining, indicating abundant polysaccharide accumulation (Fig. 4A). Conversely, the plasmidless L2R strain, which natively lacks the machinery for glycogen synthesis, demonstrated a marked reduction in PAS staining intensity. In the control strain, L2-BsciDng-dCas12-*incA*g, inclusions from both the induced and uninduced coverslips maintained dense, bright PAS staining, confirming that dCas12 expression alone does not alter glycogen accumulation. However, target knockdown in the L2-BsciDng-dCas12-*euo*g strain yielded a distinct phenotype: while the uninduced inclusions retained strong PAS reactivity, induction of dCas12 and subsequent loss of *euo* gene expression resulted in significantly diminished PAS staining intensity (Fig. 4B). Together, these data suggest that *euo* expression is required for normal glycogen accumulation within the chlamydial inclusion.

**Figure 4:**
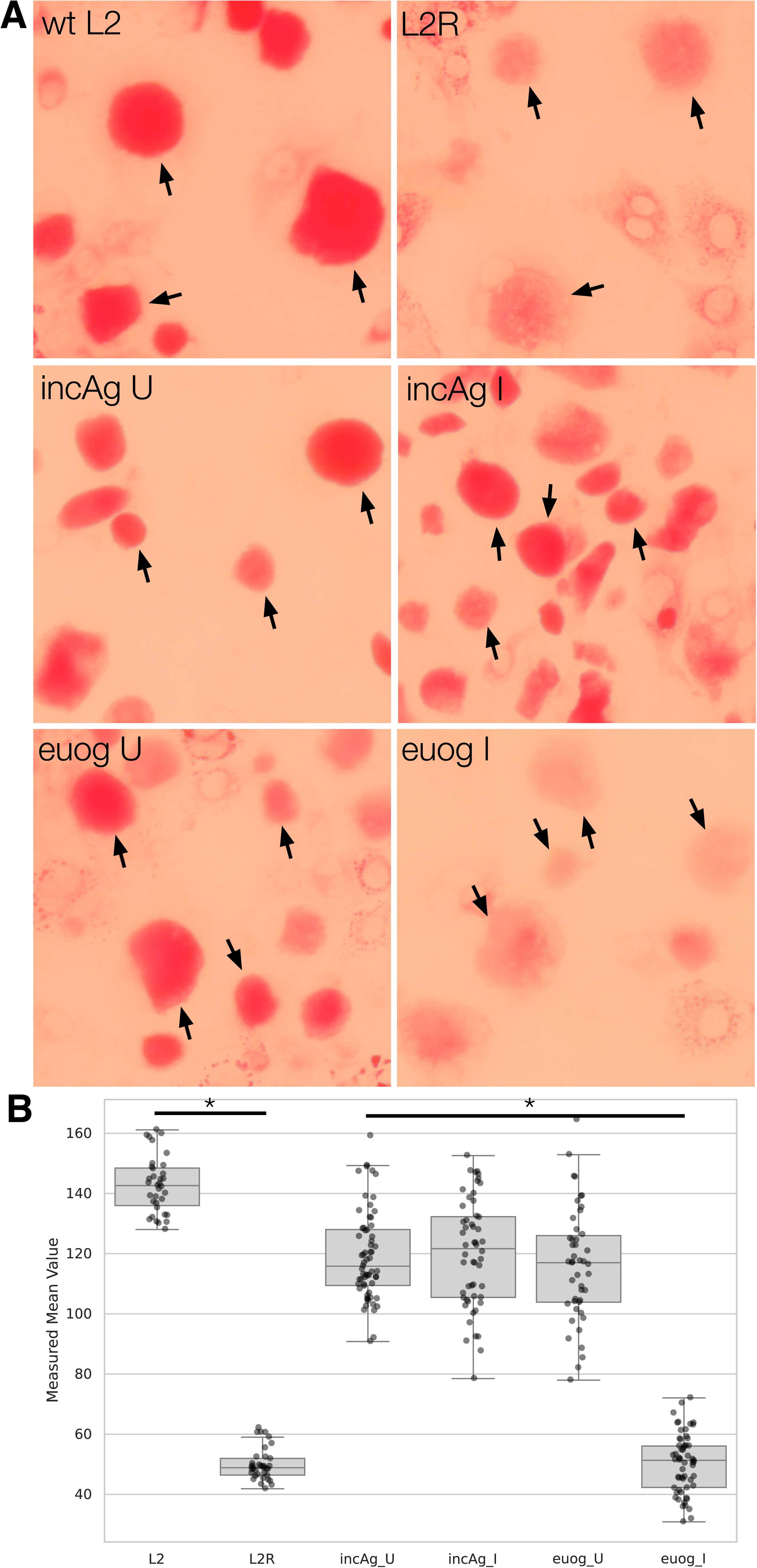
Knockdown of *euo* reduces glycogen accumulation within the chlamydial inclusion. (A) Representative fields of view of infected Cos7 cells stained with Periodic acid– Schiff (PAS) to visualize intrainclusion glycogen. Wild-type (wt) *C. trachomatis* L2 inclusions exhibit robust PAS reactivity, whereas plasmidless L2 (L2R) infections display a marked reduction in carbohydrate staining. Control infections with L2-BsciDng-dCas12-*incA*g maintain high glycogen levels regardless of induction status. Conversely, an induction of dCas12 in the strain targeting *euo* (L2-BsciDng-dCas12-*euo*g) significantly diminishes PAS staining intensity compared to its uninduced control. (B) Quantification of PAS staining intensity from at least 40 inclusions per experimental group. Data represent PAS staining from individual chlamydial inclusions. *, *(p < 0.01)*.

### Reinfectivity of *euo* knockdown EBs

Although *euo* knockdown had minimal effects on progression through the primary developmental cycle, including the formation of EB-like cells, reinfection assays revealed a marked reduction in infectious progeny. To determine the stage at which EBs produced under *euo* deficiency fail during secondary infection, we examined early events associated with EB entry into host cells.

EBs were purified from dCas12-induced or uninduced primary infections and quantified by ddPCR. Equal numbers of purified L2-BsciDng-dCas12-*euo*g EBs were used to infect fresh monolayers at an MOI of approximately 15. We then assessed translocation and phosphorylation of TarP, a type III secretion system (T3SS) effector that is delivered into host cells within minutes of EB contact and is important for efficient entry (5, 26). TarP phosphorylation was detected using an anti-phosphotyrosine antibody (4G10). Infected cells were fixed at 6 hpi and imaged by confocal microscopy.

In cells infected with EBs derived from uninduced cultures, nascent chlamydial forms were clustered near the nucleus and were positive for 4G10 staining, consistent with efficient TarP translocation and phosphorylation (Fig. 5). These organisms exhibited weak *incD*prom-driven fluorescence and lacked detectable *hctB*prom-driven scarlet-I expression, consistent with early stages of EB-to-RB differentiation (Fig. 5).

**Figure 5:**
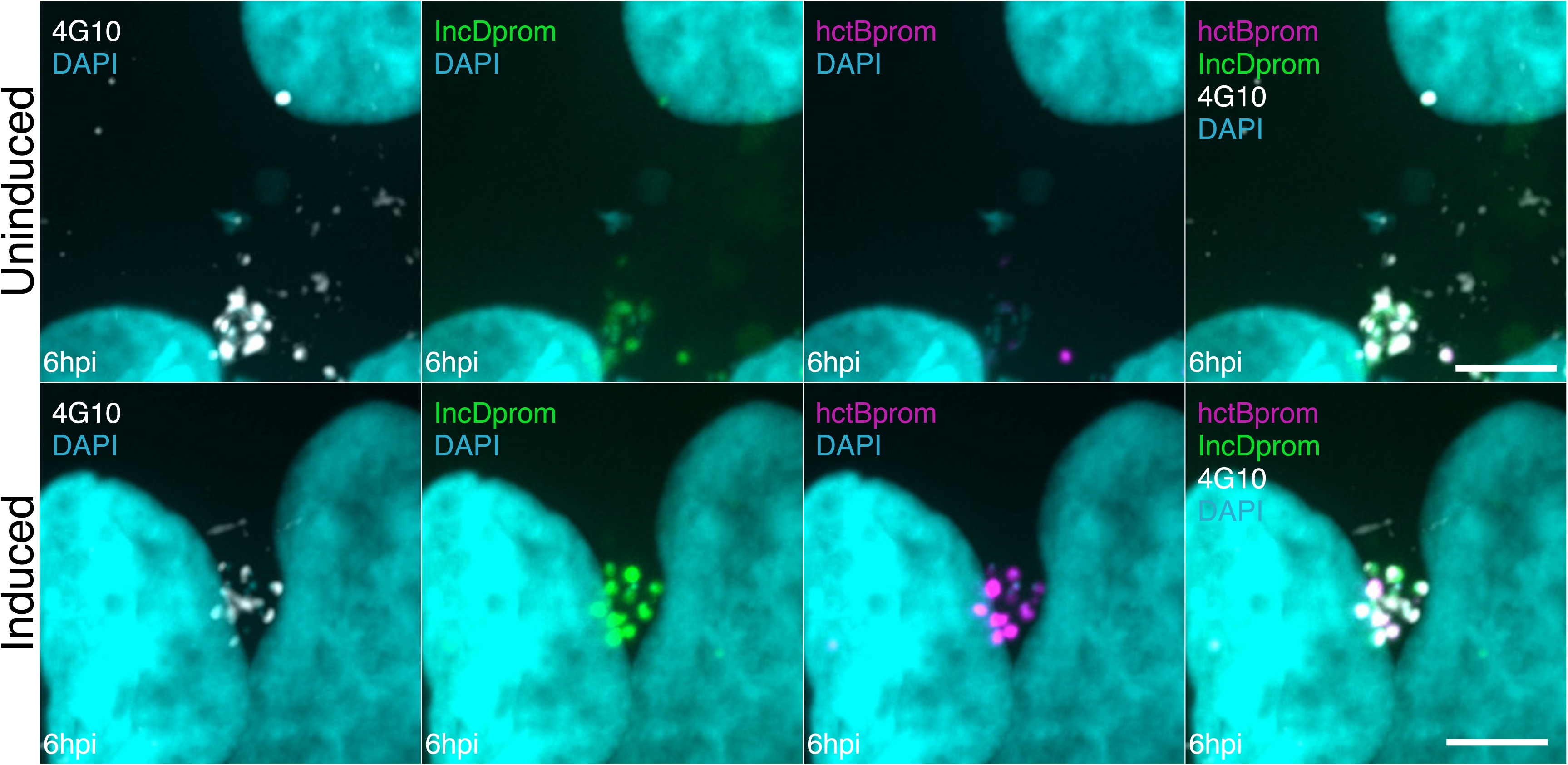
*euo* knockdown progeny exhibit normal host cell entry and TarP translocation. Representative confocal micrographs of Cos-7 cells infected with purified EBs derived from uninduced or pre-induced (*euo* knockdown) L2-BsciDng-dCas12-*euo*g primary infections. Cultures were fixed at 6 hpi to assess early infection events. Nascent chlamydial forms are identified by the *incD*prom (RB; green) and *hctB*prom (EB; magenta) reporters. Host cell entry and Type III Secretion System (T3SS) activity were visualized via immunofluorescence staining with 4G10 (white) to detect phosphorylated tyrosine residues on the translocated effector TarP. DNA was counterstained with DAPI (cyan). Perinuclear clustering and robust TarP phosphorylation in both conditions indicate that *euo* knockdown does not impair EB attachment or internalization. Scale bars = 5 µm.

Similarly, cells infected with EBs purified from *euo* knockdown cultures showed perinuclear clustering of nascent chlamydial forms that were also positive for 4G10 staining, indicating that TarP translocation and phosphorylation occurred normally (Fig. 5). However, in contrast to controls, these 4G10-positive organisms displayed markedly increased *incD*prom-driven fluorescence and robust scarlet-I expression from the *hctB*prom (Fig. 5). Notably, in wild-type infections, *hctB* promoter activity is restricted to the EB cell form and is not normally detected until approximately 20 hpi.

### Aborted development of the secondary infection

Reinfection with EBs produced under *euo* knockdown conditions resulted in a pronounced defect in establishing a productive secondary infection. This defect was not absolute, as a subset of infections progressed to form inclusions that appeared morphologically similar to those produced by wild-type bacteria. To determine the stage at which development failed, we quantified the progression of individual EBs through early stages of secondary infection.

Host cell monolayers were infected at an MOI of approximately 0.3 with EBs purified from induced or uninduced primary infections and fixed at 6, 12, and 18 hpi. Using confocal microscopy, we classified intracellular chlamydial forms as single cells (prior to division), early inclusions containing 3–10 cells (corresponding to 2–4 divisions), or fully developed inclusions containing more than 30 cells (>4 divisions).

At 6 hpi, internalized chlamydial forms from both conditions were predominantly present as single cells (Fig. 6A,B). By 12 hpi, many of the chlamydial cells derived from uninduced cultures had initiated replication, with 45% progressing to either early or fully developed inclusions (Fig. 6A,B). In contrast, only 6% of chlamydial cells derived from *euo* knockdown cultures had progressed beyond the single-cell stage at this time point. By 18 hpi, 80% of the chlamydial cells from uninduced cultures had formed fully developed inclusions, whereas only ∼25% of infections initiated with *euo* knockdown–derived EBs progressed beyond the single-cell stage, and only ∼15% formed fully mature inclusions (Fig. 6A,B).

**Figure 6:**
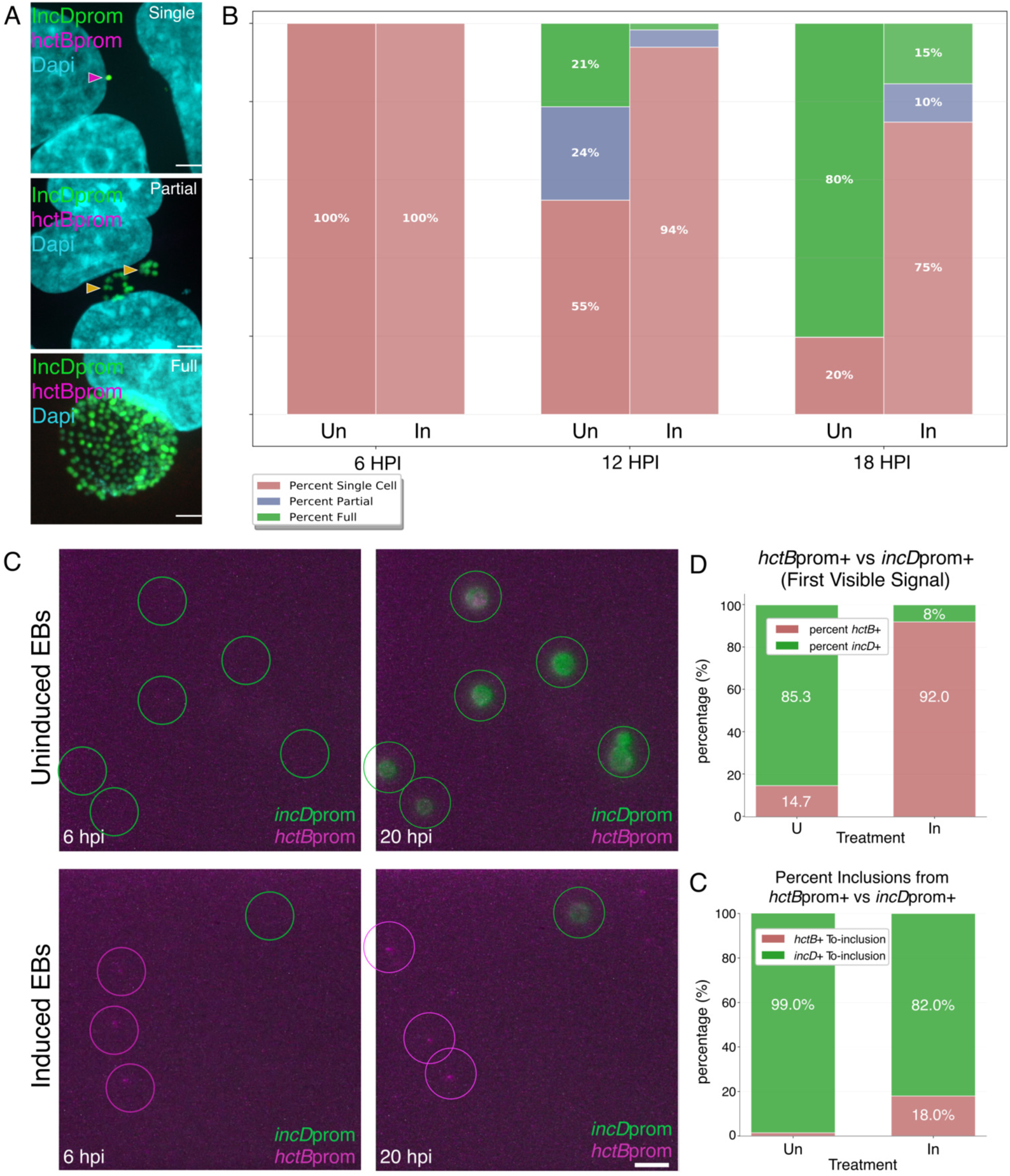
Progeny from *euo* knockdown infections exhibit a lethal germination defect during secondary infection. A) Secondary infection progression. Representative confocal micrographs of Cos-7 cells reinfected with purified EBs derived from uninduced or *euo* knockdown (induced) primary infections. Samples were fixed at 6, 12, and 18 hpi and stained with DAPI (blue). Inclusions were staged based on chlamydial density: Single Cells (pre-division), Partial (2–5 divisions), and Full (>5 divisions). Scale bar = 3µm B) Quantification of developmental arrest. Percentage of chlamydial forms at each developmental stage over the first 18 hours of secondary infection. Single cells are shown in pink, partial inclusions in purple, and full inclusions in green. Data represent the mean of 6 FOVs per treatment (n>20 inclusions per FOV). C) Representative germination dynamics. Live-cell microscopy stills at 6 and 20 hpi of individual infections initiated with uninduced or *euo* knockdown-derived EBs. Circles denote the initial detected fluorescent signal: *incD*prom (RB-initiation; green) or *hctB*prom (premature EB-regulon activation; magenta). Scale bar = 10µm. D) Temporal sequence of initial promoter activation. Percentage of inclusions categorized by the first detected fluorescent signal. *euo* knockdown progeny predominantly activated the late-stage *hctB*prom (red) prior to or instead of the RB-specific *incD*prom (green). Data represent 16 FOVs per treatment (n>5 infection events per FOV). E) Correlation between germination pattern and inclusion success. Percentage of successful versus aborted inclusions categorized by the identity of the first visible promoter signal (*incD*prom in green or hctBprom in red). Premature *hctB* activation is strongly associated with the failure to establish a productive secondary infection.

To determine whether aberrant early activation of *hctB*prom was associated with failure of germination, we used live-cell imaging to examine early gene expression dynamics in individual reinfections. Monolayers were infected at an MOI of approximately 0.3 with EBs purified from L2-BsciDng-dCas12-*euo*g cultures that were either induced or uninduced for dCas12 expression, and imaged continuously for 48 h. Time-lapse imaging allowed retrospective determination of infection outcome and early promoter activation patterns for more than 50 individual infection events per condition (Example full movies SM1 and SM2).

During early infection, we scored whether nascent germinating EBs first activated the *incD*prom or the *hctB*prom. Among germinating EBs derived from uninduced cultures, 85.3% activated *incD*prom and did not activate the *hctB*prom (Fig. 6C,D). In contrast, only 8% of germinating EBs derived from *euo* knockdown cultures exhibited initial activation of *incD*prom; instead, the majority had noticeable early *hctB*prom activity.

We next examined which early gene expression pattern was associated with productive inclusion formation. For EBs derived from uninduced cultures, 99% of fully developed inclusions originated from germinating EBs that activated *incD*prom first (Fig. 6E,F). Similarly, for EBs derived from Euo knockdown cultures, 82% of fully developed inclusions arose from germinating EBs that activated *incD*prom and had little discernible *hctB*prom activity (Fig. 6E,F). Together, these data indicate that premature or aberrant activation of *hctB*prom in EBs produced under *euo* knockdown conditions is strongly associated with aborted secondary infection.

Because residual dCas12 activity could, in principle, persist into the secondary infection and continue to repress *euo*, we tested this possibility experimentally. We generated a strain in which the *neongreen* (*ng*) gene in the reporter construct is targeted by dCas12 (L2-Eng-dCas12- *ng*g). Monolayers were infected with L2-Eng-dCas12-*ng*g and either induced with aTc at infection or left uninduced. Parallel infections were processed for either EB purification or confocal imaging. Confocal microscopy revealed strong neongreen fluorescence in inclusions from uninduced infections, whereas neongreen signal was undetectable in induced samples, confirming efficient dCas12-mediated repression during the primary infection (Fig. S2A).

EBs purified from induced and uninduced infections were then used to reinfect fresh monolayers, which were subjected to live-cell time-lapse imaging at 30-minute intervals for 36 h. Neongreen fluorescence intensity was quantified for individual inclusions (n > 20 per condition), and average expression profiles were plotted to assess *euo* promoter reactivation during the secondary infection (Fig. S2B). No differences were observed in either the timing or magnitude of neongreen reexpression between infections initiated with EBs derived from induced or uninduced primary infections. Together, these data indicate that dCas12-mediated repression during the primary infection does not persist into the secondary infection and support the conclusion that the reinfection phenotypes are not attributable to carryover of dCas12 activity.

### Late/infectivity gene expression is dysregulated during germination of EBs from *euo* knockdown infections

Gene expression during the chlamydial developmental cycle is organized into three major regulons corresponding to the RB, IB, and EB cell forms (18, 23). In the experiments described above, we observed inappropriate activation of the EB-associated *hctB* promoter during germination of EBs derived from *euo* knockdown infections. To determine which gene regulon(s) are dysregulated during this stage, we performed FISH to assess expression of representative genes from each of the three regulons.

EBs were purified from primary infections that were either induced or uninduced for dCas12-mediated *euo* knockdown. Monolayers were then infected at an MOI of approximately 15 and fixed at 8 hpi, a time point that corresponds to post-germination but precedes the first division event, which occurs at approximately 10-12 hpi. FISH was performed using probes targeting two RB-associated genes (*euo* and *incD*), one IB-associated gene (*hctA*), and three EB-associated genes (*hctB, incM*, and *incV*). Confocal images were acquired from 10 randomly selected infected cells per condition, and FISH signal intensity was quantified for intracellular chlamydial cells.

Germinating *Chlamydia* derived from uninduced cultures showed little to no detectable signal for the EB-associated genes *hctB*, *incM*, and *incV* (Fig. 7A–C). Similarly, no detectable signal was observed for the IB-associated gene *hctA* (Fig. 7D). In contrast, robust mRNA signals for the RB-associated genes *euo* and *incD* were readily detected, consistent with normal EB-to-RB transition during germination (Fig. 7E,F).

**Figure 7:**
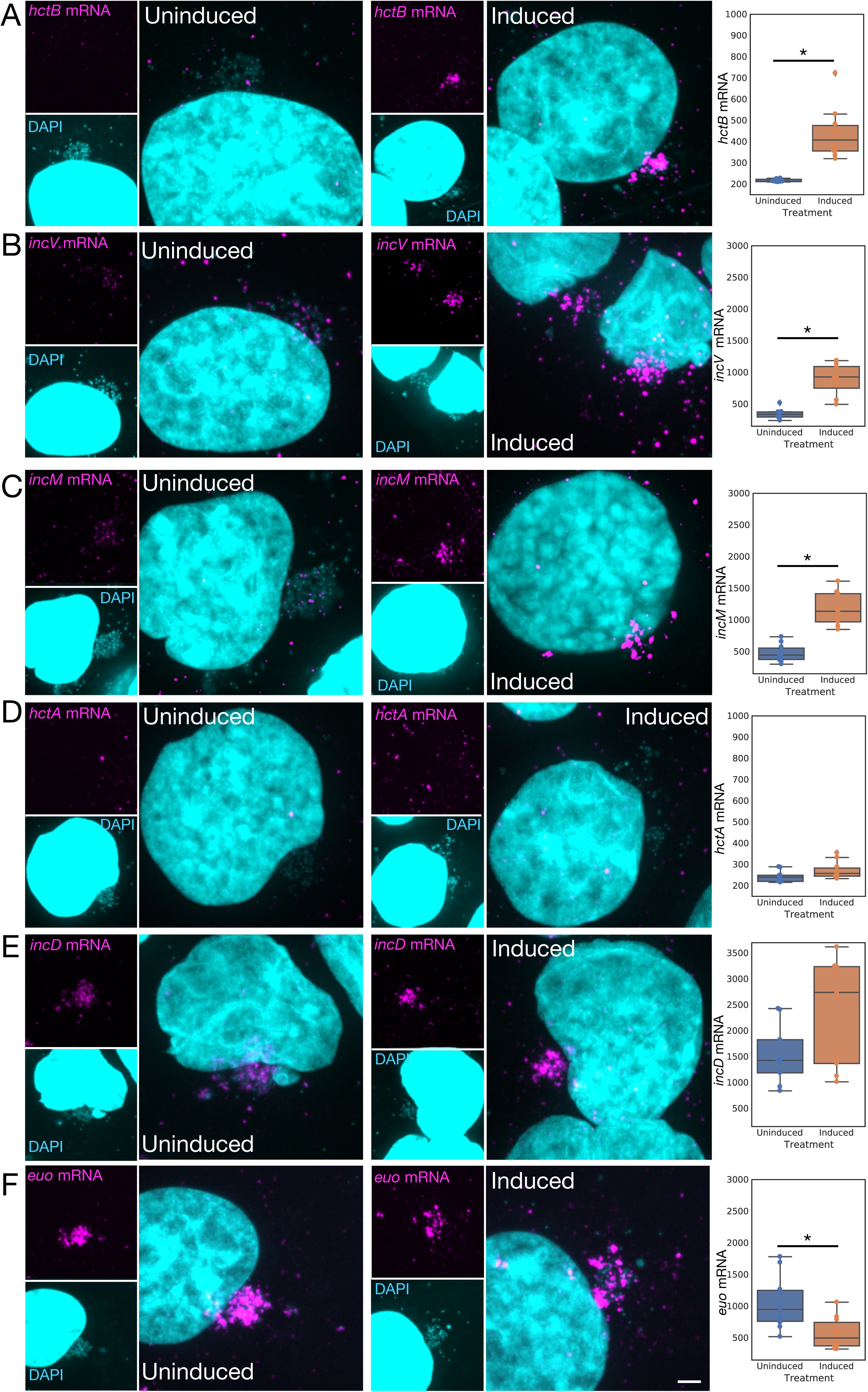
Knockdown of *euo* results in premature and specific dysregulation of the EB regulon during germination. Representative confocal micrographs and quantitative FISH analysis of Cos-7 cells infected with purified EBs from uninduced or *euo* knockdown (induced) primary infections. Samples were fixed at 8 hpi to capture the transcriptional state of germinating chlamydial cells prior to the first division. A-F) Monolayers were processed for FISH using probes (magenta) targeting transcripts from the three major developmental regulons: RB-associated (*euo*, *incD*; panels E and F), IB-associated (*hctA*; panel D), and EB-associated (*hctB*, *incV, incM*; panels A, B and C). DNA was counterstained with DAPI (cyan). Quantification (right): Fluorescence intensity was measured per chlamydial cluster to determine transcript abundance. While RB-associated transcripts were detected in both conditions, *euo* knockdown progeny exhibited significant and inappropriate activation of the EB-specific regulon genes (*incM, incV, hctB*). No activation of the IB-associated *hctA* was observed in either condition. Data represent the mean signal intensity of n>10 clusters per treatment. Scale bar = 4 µm. * = p<0.01 (Student’s t-test).

In contrast, germinating *Chlamydia* derived from *euo* knockdown infections exhibited strong mRNA signals for all three EB-associated genes (*hctB*, *incM*, and *incV*) (Fig. 7A–C). Notably, expression of the IB-associated gene *hctA* remained undetectable (Fig. 7D), indicating that dysregulation was specific to the EB regulon. Signals for the RB-associated genes *incD* and *euo* were also detected in *euo* knockdown–derived *Chlamydia*; however, *euo* signal intensity was mildly reduced relative to uninduced controls, while *incD* levels were slightly increased but more variable (Fig. 7E,F).

Together, these data indicate that *euo* knockdown during a primary infection leads to premature and inappropriate activation of the EB/infectivity regulon during germination upon reinfection, a defect that correlates with failure to establish a productive secondary infection.

## Discussion

The chlamydial infectious cycle is an intracellular developmental process involving three distinct morphologic forms: the Elementary Body (EB), the Intermediate Body (IB), and the Reticulate Body (RB). This cycle can be described by three corresponding regulons, which are defined by the expression of specific DNA-binding proteins: Euo (RB-specific), HctA (IB-specific), and HctB (EB-specific) (11, 12, 18).

The infection begins when the EB, a metabolically quiescent form, triggers its own entry into the host cell via pathogen-directed endocytosis (5, 26, 27). Once internalized, the EB initiates a gene expression cascade that facilitates its differentiation into the replication-competent RB. Following several rounds of division, these RBs begin to produce IBs which subsequently undergo a final maturation phase, lasting approximately 8–10 hours, to regenerate the infectious EB, thereby completing the cycle (11, 12).

Previous studies showed that Euo is a DNA binding protein/transcriptional regulator that is proposed to act in repressing late cycle genes (both IB and EB) (14, 15). Additionally, we have shown that ectopic expression of Euo resulted in a reversible arrest of the development cycle. The arrested chlamydial cells were trapped phenotypically at an early IB stage of the cycle. These cells had halted DNA replication initiation but had not shifted gene expression from RB like to IB/EB like. This arrested state was dependent on continued expression of Euo. When ectopic expression was halted, native *euo* expression was repressed and the cycle resumed (16). While Euo is highly conserved across the *Chlamydiota* phylum, it has no known orthologs in other bacterial lineages, suggesting a unique and essential role in chlamydial biology. Currently, its mechanistic function in developmental transitions has remained elusive.

### Euo is Dispensable for Primary Developmental Kinetics and Replication

In the present study, we utilized CRISPRi-mediated knockdown to determine if Euo is required for progression through specific stages of the developmental cycle. Unexpectedly, although Euo depletion dramatically reduced the production of infectious progeny, loss of Euo expression did not alter the temporal dynamics of the primary infection. Live-cell imaging using RB-, IB-, and EB-specific reporters (*incD*prom, *porB*prom, and *hctB*prom) showed that the sequence and timing of developmental transitions remained intact (Fig. 2A). Furthermore, the lack of impact on chromosomal replication and the index of replication (iRep) suggests that Euo is not involved in the population-level shift from replication (RB) to differentiation (IB, EB) (Fig. 2B, C). This is consistent with our observation that individual cell morphologies—including the ultrastructure of condensed EBs—appear normal under TEM and confocal microscopy (Fig. 3C).

Notably, we observed a significant alteration in the spatial organization of the chlamydial IBs, which formed large clustered aggregates rather than discrete entities (Fig. 3A, B). This phenotype resembled the aberrant spatial arrangement previously reported in *Chlamydia* strains defective in intrainclusion glycogen production (20, 28, 29). Periodic Acid–Schiff (PAS) staining confirmed that *euo* knockdown significantly reduced glycogen accumulation within the inclusion lumen. While the precise mechanistic basis for this metabolic defect remains unknown, Euo is known to physically interact with the plasmid-encoded regulator Pgp4 (30). Pgp4 is a plasmid-encoded protein that acts in the regulation of genes involved in glycogen synthesis and loss of pgp4 results in a reduction of glycogen in the inclusion (23). It is therefore intriguing to speculate that the Euo–Pgp4 regulatory axis plays a role in inter-inclusion glycogen accumulation.

### Euo establishes the EB transcriptional state required for productive germination

Although the primary developmental cycle proceeded with normal kinetics, the EBs produced under *euo* knockdown conditions were markedly impaired in their ability to initiate a secondary infection. This defect did not arise during attachment, entry, or early type III secretion, as TarP phosphorylation occurred at levels comparable to controls (Fig. 5). Instead, the block emerged during EB germination.

During a typical infection, germinating EBs initiate a tightly ordered transcriptional program in which RB-specific genes (e.g., *incD, euo*) are activated first, enabling the transition to a replication-competent RB. In contrast, EBs generated under Euo-depleted conditions displayed a pronounced dysregulated transcriptional phenotype upon re-infection of a cell monolayer. These EBs simultaneously activated RB-associated genes while aberrantly re-expressing EB/infectivity-associated genes, including *hctB, incM*, and *incV* (Fig. 7). This inappropriate reactivation of late-stage genes strongly correlated with developmental failure, as only the minority of EBs that followed the canonical *incD*-first activation pattern progressed to form productive inclusions (Fig. 6E, F).

Despite this defect, Euo-depleted EBs had normal, condensed nucleoids, indicating that gross EB morphogenesis remained intact. The selective misregulation of EB-associated genes— without concomitant activation of IB-associated markers such as *hctA*—points to a specific failure in transcriptional control during the EB-to-RB transition rather than a global developmental disruption. Importantly, we demonstrated that dCas12 activity does not persist into the secondary infection (Fig. S2), and *euo* transcripts were readily detected during germination. Together, these observations indicate that the observed phenotype reflects defective transcriptional programming of the EB during its formation in the primary infection, underscoring a critical role for Euo in establishing the EB transcriptional state necessary for successful germination.

## Conclusion

In summary, Euo expression does not appear to control a switch for primary RB-to-EB transition, but it appears to be a critical regulator of the EB-RB germination step of the chlamydial life cycle. Our results support a novel role for Euo in creating or maintaining the EB infectious-state. We propose that EB germination exists in a biphasic state: if the EB/late-gene regulon is initiated first, development is aborted; however, if the RB regulon is established first, the infection proceeds normally. Our findings indicate that Euo is a critical factor in biasing this competition toward RB gene expression. Interestingly, once the RB regulon is established, continued Euo expression appears dispensable.

While our previous work showed that overexpression of Euo arrests development by preventing RB-to-EB maturation, the knockdown phenotype observed here resulted in only minor spatial defects during the primary cycle. This suggests that while high levels of Euo can block the transition to late-cycle forms, endogenous levels are primarily required to ensure the transcriptional fidelity of the resulting progeny. Future studies will investigate whether Euo interacts or works in concert with other global regulators, such as HctA, HctB or other regulatory proteins, to stabilize the condensed, transcriptionally silent state of the infectious EB. We hypothesize that DNA binding proteins in the EB ensure a productive gene expression cascade upon germination.

## Materials and Methods

### Cell Culture

Cos-7 cells obtained from ATCC were maintained using RPMI 1640 supplemented with 10% Fetalplex and treated with 10 mg/mL gentamicin to prevent contamination. Cultures were grown at 37°C with 5% CO_2_. All *C. trachomatis* L2-bu434 (L2) infections were carried out in Cos-7 cells. EBs isolated from infectious cultures were harvested and purified utilizing centrifugation over a 30% MD-76R density gradient. Purified EBs were stored in sucrose-phosphate-glutamate buffer (10 mM sodium phosphate [8 mM K2HPO4, 2 mM KH2PO4], 220 mM sucrose, and 0.50 mM L-glutamic acid, pH 7.4) at −80°C.

### Vector Construction

All constructs used p2TK2-SW2 (31) as the backbone, and cloning was performed using the In-fusion HD EcoDry Cloning kit (Thermo Fisher Scientific, Waltham, MA, USA). Primers were ordered from Integrated DNA Technologies (IDT, Coralville, IA, USA) and noted in supplementary table S1.

To create the Scarlet-I reporter cassettes, *hctB*prom_Scarlet-*incD*prom_neongreen (BsciDng) and *porB*prom_Scarlet-*incD*prom_neongreen (PsciDng), *incD*prom was amplified from L2 genomic and used to replace *euo*prom in the BsciEng and PsciEng reporter cassettes described previously (18).

CRISPRi constructs utilized the previously described dCas12 system cloned into our BsciDng and PsciDng reporter plasmids (32, 33). Guide RNA for *incA* and *euo* promoters as well as primers utilized are indicated in supplemental table 1.

### Live-cell Microscopy

Cos-7 monolayers were grown on glass bottom 6-well plates, and temperature and CO_2_ were maintained using an OKOtouch stage incubator. A Nikon Eclipse TE300 inverted microscope was used for live cell imaging using a 20X, 0.4-numeric-aperture objective or a 60X 1.4 NA objective depending on experiment. Fluorescent protein excitation was achieved using a ScopeLED lamp at 470 and 595 nm along with BrightLine bandpass filters at 514/30 and 590/20 nm. DIC was used for focusing. Image acquisition was achieved using an Andor Zyla sCMOS camera. Micro-Manager software was used to image every 30 minutes. Experiment data analysis utilized matplotlib, pandas, and seaborn using custom Python notebooks to determine and utilize Trackmate software to measure fluorescence intensities in individual inclusions over time (12).

### Chlamydial Transformation

*C. trachomatis* transformation was performed as previously described using 500 ng/μL spectinomycin for selection (12, 34). Clonal populations were generated by inclusion isolation with a micromanipulator. The plasmids from the chlamydial transformants were verified by sequencing.

### Infections

Cos-7 cell monolayers were incubated with infectious EBs in Hanks Balanced Salt Solution (HBSS) (Gibco) for 15 minutes at 37°C. The inoculum was then removed, and the host cells were washed with pre-warmed HBSS. Infections with pre-treated (uninduced and induced) EBs were additionally washed 2 x 5min with pre-warmed HBSS + 1mg/mL heparin to remove unbound EBs. After washing, the HBSS was replaced with fresh RPMI 1640 containing 10% fetal bovine serum, 10 µg/mL gentamicin, 1 µg/mL cycloheximide, and 1 mg/mL heparin to ensure synchronization of infection and to lower reinfection in long experiments.

### Confocal Microscopy

Infected monolayers were fixed overnight in 2% paraformaldehyde at 4°C. They were washed the next day with phosphate-buffered saline and stained with DAPI to visualize DNA. Coverslips were mounted using MOWIOL mounting solution (100 mg/mL MOWIOL 4-88, 25% glycerol, 0.1 M Tris, pH 8.5). Confocal images were acquired using a Nikon CrestOptics X-Light confocal coupled with Nikon Elements imaging software. Images were taken using a 100× oil-immersion objective. Monoclonal antibody 4G10 (Millipore) at a dilution of 1:200 was used to detect phosphotyrosine.

### Transmission Electron Microscopy

For analysis of the structure of Ctr post knockdown, cell monolayers were infected with the indicated strain at an MOI of > 1 and induced with 3ng aTc at infection. Infected cells were harvested via sonication at 48 hpi and purified utilizing centrifugation over a 30% MD-76R density gradient. Pellets were rinsed with 1× phosphate-buffered saline (PBS), and the pellet was fixed with EM fixative (2% paraformaldehyde [PFA], 2% glutaraldehyde, 0.1M phosphate buffer, pH 7.2) overnight at 4°C. Fixed pellets were post-fixed with 1% osmium tetroxide, rinsed and dehydrated before embedding with Spurr’s resin and cross-sectioned with an ultramicrotome (Reichert Ultracut R; Leica). Ultra-thin sections were placed on formvar-coated slot grids and stained with uranyl acetate and Reynolds lead citrate. TEM imaging was conducted with a Tecnai G2 transmission electron microscope (FEI Company; Hillsboro, OR).

### Replating Assay

Ctr were harvested via sonication at 48 hpi and purified utilizing centrifugation over a 30% MD-76R density gradient. Infectious EBs were then replated in a 96-well plate (normalized to chromosomes/µl) on fresh Cos-7 monolayers in a twofold dilution series. Re-infected monolayers were grown for 30 hpi prior to fixation with methanol before being stained with DAPI and Ctr MOMP polyclonal antibody, FITC (Fishersci). DAPI was used for visualization of host nuclei. Anti-Ctr antibody was used for staining of re-infected inclusions for IFU counts. Inclusions were imaged using a Nikon Eclipse TE300 inverted microscope, utilizing a scopeLED lamp at 470 and 390 nm, and BrightLine band pass emissions filters at 514/30 and 434/17 nm. An Andor Zyla sCMOS camera was used in conjunction with Micro-Manager software for image acquisition.

### Digital Droplet PCR

Ctr genomes were isolated using an Invitrogen PureLink Genomic DNA Minikit. Samples were then diluted. DNA samples were added to ddPCR Supermix (Bio-RAD). Amplification was achieved using primer/probe sets for nqrA (origin) or pyrG (terminus). Droplets were generated using a Bio-RAD: QX200 AutoDG Droplet Digital PCR System. Data analysis was performed using Bio-RAD QX Manager 1.2 along with custom matplotlib, pandas, and seaborn Python scripts.

### Periodic Acid–Schiff (PAS) Staining

Ctr-infected monolayers were fixed with ice cold 100% methanol for 10 minutes, followed by three washes with PBS. To detect chlamydial glycogen-containing inclusions, coverslips were treated with 1% periodic acid (Fishersci) for 5 minutes to oxidize the carbohydrates. After rinsing thoroughly with distilled water, the cells were incubated in Schiff’s reagent (Fishersci) for 15 minutes at room temperature in the dark to develop the staining. The coverslips were washed in running tap water for 5 minutes to intensify the color. Samples were visualized and imaged using brightfield microscopy.

### Fluorescent In-situ Hybridization

Ctr-infected monolayers were fixed with 4% PFA for 10 minutes at room temperature. Fixed coverslips were then permeabilized in 70% ethanol at −20°C overnight. Hairpin amplification was achieved using Molecular Instruments HCR FISH Kit. The custom-designed probes were ordered from Molecular Instruments and are listed in Table S1. Coverslips were mounted using MOWIOL and imaged using the CrestOptics X-Light confocal system.

Quantification of the FISH signal was performed using imageJ (FIJI, https://imagej.net). Confocal Z stack images were z projected using sum. The perinuclear migrated inclusions were identified in the DAPI channel and ROI was created around them. The FISH signal intensity was measured and the mean intensity was reported.

---

**Figure S1. Schematic representation of cell form reporter CRISPRi plasmid constructs.**

Dual-promoter reporter cassettes were engineered into the p2tK2_shuttle vector backbone alongside a CRISPRi-mediated gene knockdown system. The CRISPRi system contains specific guide RNAs (gRNAs) targeting either *euo* or *incA* open reading frames. The resulting recombinant plasmids were successfully transformed into *Chlamydia trachomatis* (Ctr).

**Figure S2. CRSIPRi knockdown does not persist into the secondary infection.**

Parallel monolayers were infected with L2-Eng-dCas12-*ng*g. (A) One set of cultures was fixed and imaged to assess neongreen expression. Bright neongreen–positive inclusions were observed exclusively in uninduced cultures, confirming efficient knockdown upon induction. (B) At 48 hpi, EBs were harvested from induced and uninduced infections and used to infect fresh monolayers grown on glass-bottom plates. Secondary infections were imaged every 30 minutes for 40 hours. Neongreen fluorescence intensity was quantified for >20 individual inclusions, and the mean (solid lines) with standard error (shaded regions) is shown. Neongreen expression was indistinguishable between infections initiated with EBs derived from induced versus uninduced primary cultures, indicating that dCas12-mediated knockdown did not carry over into the secondary infection. Scale bar = 20 µm.

**Supplemental Movie MS1. Microscopic analysis of nascent chlamydial germination during CRISPRi-mediated *euo* knockdown.** Confluent Cos-7 monolayers were infected with *Chlamydia trachomatis* L2-BsciDng-dCas12-*euo*g elementary bodies (EBs) under conditions inducing *euo* knockdown. Live-cell imaging using a 60X 1.4 NA objective was performed at 30-minute intervals initiated immediately post-infection to track the differential expression of the EB-specific hctBp-mScarlet-I reporter and the reticulate body (RB)-specific incDprom-mNeonGreen reporter. Nascent intracellular forms that aberrantly activated the hctB promoter prior to euo promoter expression failed to differentiate into productive inclusions. Conversely, productive inclusions originated exclusively from germinating forms that expressed mNeonGreen while remaining negative for mScarlet-I during early developmental stages.

**Supplemental Movie MS2. Microscopic analysis of nascent chlamydial germination under non-inducing control conditions.** Confluent Cos-7 monolayers were infected with non-induced *Chlamydia trachomatis* L2-BsciDng-dCas12-*euo*g elementary bodies (EBs) serving as the experimental control. Live-cell imaging using a 60X 1.4 NA objective was performed at 30-minute intervals immediately post-infection to monitor reporter expression dynamics. Under these non-knockdown conditions, germinating forms consistently avoided aberrant, early activation of the hctB promoter, exclusively expressing the *euo* promoter-driven neongreen reporter during early developmental stages to yield productive inclusions.

## References

1. Rockey DD, Matsumoto A. 2000. The chlamydial developmental cycle. Prokaryotic Dev 403–425.

2. Abdelrahman YM, Belland RJ. 2005. The chlamydial developmental cycle. FEMS Microbiol Rev 29:949–59.

3. Clifton DR, Dooley CA, Grieshaber SS, Carabeo RA, Fields KA, Hackstadt T. 2005. Tyrosine phosphorylation of the chlamydial effector protein Tarp is species specific and not required for recruitment of actin. Infect Immun 73:3860–8.

4. Chen Y-S, Bastidas RJ, Saka HA, Carpenter VK, Richards KL, Plano GV, Valdivia RH. 2014. The Chlamydia trachomatis type III secretion chaperone Slc1 engages multiple early effectors, including TepP, a tyrosine-phosphorylated protein required for the recruitment of CrkI-II to nascent inclusions and innate immune signaling. PLoS Pathog 10.

5. Romero MD, Carabeo RA. 2024. Dynamin-dependent entry of Chlamydia trachomatis is sequentially regulated by the effectors TarP and TmeA. Nat Commun 15:4926.

6. Scidmore MA, Fischer ER, Hackstadt T. Restricted fusion of Chlamydia trachomatis vesicles with endocytic compartments during the initial stages of infection. Infect Immun 71:973–984.

7. Grieshaber SS, Grieshaber NA, Hackstadt T. 2003. Chlamydia trachomatis uses host cell dynein to traffic to the microtubule-organizing center in a p50 dynamitin-independent process. J Cell Sci 116:3793–802.

8. Elwell CA, Jiang S, Kim JH, Lee A, Wittmann T, Hanada K, Melancon P, Engel JN, Valdivia RH. 2013. Correction: Chlamydia trachomatis Co-opts GBF1 and CERT to Acquire Host Sphingomyelin for Distinct Roles during Intracellular Development. PLoS Pathog 9.

9. Derré I. Chlamydiae interaction with the endoplasmic reticulum: contact, function and consequences. Cell Microbiol 17:959–966.

10. Abdelrahman Y, Ouellette SP, Belland RJ, Cox JV. 2016. Polarized Cell Division of Chlamydia trachomatis. PLoS Pathog 12.

11. Chiarelli TJ, Grieshaber NA, Appa CR, Grieshaber SS. 2023. Computational Modeling of the Chlamydial Developmental Cycle Reveals a Potential Role for Asymmetric Division. mSystems 8:e00053–23.

12. Chiarelli TJ, Grieshaber NA, Omsland A, Remien CH, Grieshaber SS. 2020. Single-Inclusion Kinetics of Chlamydia trachomatis Development. MSystems 5.

13. Domman D, Horn M. 2015. Following the Footsteps of Chlamydial Gene Regulation. Mol Biol Evol 32:3035–3046.

14. Rosario CJ, Hanson BR, Tan M. The transcriptional repressor EUO regulates both subsets of Chlamydia late genes. Mol Microbiol 94:888–897.

15. Hakiem OR, Rizvi SMA, Ramirez C, Tan M. 2023. Euo is a developmental regulator that represses late genes and activates midcycle genes in *Chlamydia trachomatis*. mBio 14:e00465–23.

16. Appa Cody R., Grieshaber Nicole A., Yang Hong, Omsland Anders, McCormick Sean, Chiarelli Travis J., Grieshaber Scott S. 2024. The chlamydial transcriptional regulator Euo is a key switch in cell form developmental progression but is not involved in the committed step to the formation of the infectious form. mSphere 0:e00437–24.

17. Zhang L, Douglas AL, Hatch TP. 1998. Characterization of a Chlamydia psittaci DNA binding protein (EUO) synthesized during the early and middle phases of the developmental cycle. Infect Immun 66:1167–1173.

18. Grieshaber NA, Appa C, Ward M, Grossman A, McCormik S, Grieshaber BS, Chiarelli T, Yang H, Omsland A, Grieshaber SS. 2025. The T3SS structural and effector genes of Chlamydia trachomatis are expressed in distinct phenotypic cell forms. Front Cell Infect Microbiol Volume 15–2025.

19. Grieshaber NA, Runac J, Turner S, Dean M, Appa C, Omsland A, Grieshaber SS. 2021. The sRNA Regulated Protein DdbA Is Involved in Development and Maintenance of the Chlamydia trachomatis EB Cell Form. Front Cell Infect Microbiol 11.

20. Carlson JH, Whitmire WM, Crane DD, Wicke L, Virtaneva K, Sturdevant DE, Kupko JJ, Porcella SF, Martinez-Orengo N, Heinzen RA, Kari L, Caldwell HD. The Chlamydia trachomatis plasmid is a transcriptional regulator of chromosomal genes and a virulence factor. Infect Immun 76:2273–2283.

21. Gong S, Yang Z, Lei L, Shen L, Zhong G. Characterization of Chlamydia trachomatis Plasmid-Encoded Open Reading Frames. J Bacteriol 195:3819–3826.

22. Song L, Carlson JH, Whitmire WM, Kari L, Virtaneva K, Sturdevant DE, Watkins H, Zhou B, Sturdevant GL, Porcella SF, McClarty G, Caldwell HD. Chlamydia trachomatis plasmid-encoded Pgp4 is a transcriptional regulator of virulence-associated genes. Infect Immun 81:636–644.

23. Grieshaber NA, Chiarelli TJ, Appa CR, Neiswanger G, Peretti K, Grieshaber SS. 2022. Translational gene expression control in Chlamydia trachomatis. PLoS ONE 17.

24. Leblond CP, Glegg RE, Eidinger D. 1957. Presence of carbohydrates with free 1,2-glycol groups in sites stained by the periodic acid-Schiff technique. J Histochem Cytochem Off J Histochem Soc 5:445– 458.

25. Manoj MG, Shelly D. 2016. Periodic acid Schiff’s: A definitive stain in histopathological diagnosis of Cylindroma. Med J Armed Forces India 72:404–406.

26. Clifton DR, Fields KA, Grieshaber SS, Dooley CA, Fischer ER, Mead DJ, Carabeo RA, Hackstadt T. 2004. A chlamydial type III translocated protein is tyrosine-phosphorylated at the site of entry and associated with recruitment of actin. Proc Natl Acad Sci U S A 101:10166–10171.

27. Jewett TJ, Miller NJ, Dooley CA, Hackstadt T. The conserved Tarp actin binding domain is important for chlamydial invasion. PLoS Pathog 6.

28. Gehre L, Gorgette O, Perrinet S, Prevost M-C, Ducatez M, Giebel AM, Nelson DE, Ball SG, Subtil A. 2016. Sequestration of host metabolism by an intracellular pathogen. eLife 5:e12552.

29. Nguyen BD, Valdivia RH. 2010. Virulence determinants in the obligate intracellular pathogen Chlamydia trachomatis revealed by forward genetic approaches. Proc Natl Acad Sci 109:1263–1268.

30. Zhang Q, Rosario CJ, Sheehan LM, Rizvi SM, Brothwell JA, He C, Tan M. 2020. The Repressor Function of the Chlamydia Late Regulator EUO Is Enhanced by the Plasmid-Encoded Protein Pgp4. J Bacteriol 202.

31. Agaisse H, Derré I. 2013. A C. trachomatis cloning vector and the generation of C. trachomatis strains expressing fluorescent proteins under the control of a C. trachomatis promoter. PLoS ONE 8.

32. Monahan Colleen C., Held Kiara, Yang Hong, Sotelo Geselle, Grieshaber Nicole, Grieshaber Scott, Omsland Anders. 2025. Ectopic overexpression and CRISPRi-based knockdown of Chlamydia trachomatis ObgE inhibits RB replication and EB reformation. J Bacteriol 0:e00282–25.

33. Ouellette SP. 2018. Feasibility of a Conditional Knockout System for Chlamydia Based on CRISPR Interference. Front Cell Infect Microbiol 8.

34. Wang Y, Kahane S, Cutcliffe LT, Skilton RJ, Lambden PR, Clarke IN. 2011. Development of a transformation system for Chlamydia trachomatis: restoration of glycogen biosynthesis by acquisition of a plasmid shuttle vector. PLoS Pathog 7.

